# Coevolution-based prediction of key allosteric residues for protein function regulation

**DOI:** 10.1101/2022.07.25.501401

**Authors:** Juan Xie, Weilin Zhang, Xiaolei Zhu, Minghua Deng, Luhua Lai

**Author notes:** Corresponding author: Luhua Lai, College of Chemistry and Molecular Engineering, Peking University, Beijing 100871, China.

## Abstract

Allostery is fundamental to many biological processes. Due to the distant regulation nature, how allosteric mutations, modifications and effector binding impact protein function is difficult to forecast. In protein engineering, remote mutations cannot be rationally designed without large scale experimental screen. Allosteric drugs have raised much attention due to their high specificity and possibility of overcoming existing drug-resistant mutations. However, optimization of allosteric compounds remains challenging. Here, we developed a novel computational method KeyAlloSite to predict allosteric site and to identify key allosteric residues (allo-residues) based on the evolutionary coupling model. We found that protein allosteric sites are strongly coupled to orthosteric site compared to non-functional sites. We further inferred key allo-residues by pairwise comparing the difference of evolutionary coupling scores of each residue in the allosteric pocket with the functional site. Our predicted key allo-residues are in accordance with previous experimental studies for typical allosteric proteins like BCR-ABL1, Tar and PDZ3, as well as key cancer mutations. We also showed that KeyAlloSite can be used to predict key allosteric residues distant from the catalytic site that are important for enzyme catalysis. Our study demonstrates that weak coevolutionary couplings contain important information of protein allosteric regulation function. KeyAlloSite can be applied in studying the evolution of protein allosteric regulation, designing and optimizing allosteric drugs, performing functional protein design and enzyme engineering.

## Introduction

Allostery commonly refers to one type of distant regulation, that is, a perturbation at one site of a macromolecule can affect the function of another site^1^, which plays important roles in many biological processes, such as enzyme catalysis^2^ and signal transduction^3^. Compared to traditional orthosteric drugs, allosteric drugs have unique advantages, including higher specificity, fewer side effects, etc^4, 5, 6^. However, optimization of allosteric molecules faces great challenges as allosteric molecules usually have flat structure-activity relationship (SAR) and higher binding affinity does not always correspond to better activity ^7, 8, 9, 10, 11^. In an allosteric pocket, the contribution of each residue to the allosteric effect is different. Bi et al. found that the interactions between allosteric molecules and the target protein can be divided into two types, interactions that only contribute to binding, and interactions that contribute to both binding and signaling^12^. Nussinov *et al*. proposed that the atoms of allosteric effectors could be divided into anchor atoms and driver atoms. The anchors docked into the allosteric pockets, which allowed the drivers to perform a “pull” or “push” action^13^. Both drivers and anchors showed specific interactions with their host proteins, with the former mainly responsible for the allosteric efficacy and the latter for binding affinity (potency)^14^. Therefore, it is important to identify residues in the host protein that form the signaling interactions with allosteric molecules, which we refer as key allo-residues, so that allosteric molecules can be optimized and designed based on these key allo-residues. Unfortunately, identifying key allo-residues remains challenging. Currently available computational methods mainly focused on the prediction of allosteric sites^15, 16^, allosteric pathways^17, 18^ and key residues in allosteric pathways^19^. Kalescky *et al*. developed the rigid residue scan method to identify key residues for protein allostery, in which multiple molecular dynamics (MD) simulations need to be performed for unbound and bound proteins. As only one residue was regarded as a single rigid body in each simulation, many simulations were necessary, which is computationally expensive and time-consuming^20^. Therefore, methods for systematically and rapidly identifying key allo-residues in protein allosteric pockets need to be developed.

During evolution, unrelated residues may evolve independently, while functionally coupled residues coevolve. Coevolution means that when a residue changes, the residues that are structurally or functionally coupled with it will also change accordingly to maintain the overall spatial structure and biological function^21^. In principle, since homologous sequences record the long-term evolution of a protein family, the coupling pattern between residues can be estimated from multiple sequence alignment^22^.

Various methods have been developed to analyze residue-residue coupling during evolution, which greatly expedite the recent progress of protein structure prediction^23, 24, 25^. Direct coupling analysis (DCA) is one of such approaches that can remove the indirect correlation between residues and reflect the direct coevolution between residues^26^. DCA mainly uses methods in statistical physics to infer the pairwise coupling *J_ij_* between positions, which can explain the observed correlation between residues in a multiple sequence alignment. In structure predictions, only the top couplings in *J_ij_* were used^23, 27^. Recent studies showed that the weak, non-contact couplings in *J_ij_* are significantly important for the prediction of protein function, although they play as noise in predicting structural contacts. Salinas *et al*. proposed that the information of allosteric energy interactions is included in the statistics of multiple sequence alignments, and therefore is part of the entire evolutionary constraints^28^. Russ *et al*. found that the top coupling items in *J_ij_* alone cannot effectively reproduce the alignment statistics in the AroQ family or the functional effects of mutations. This implies that protein functions may depend on many weak, non-contact items in *J_ij_*. Although there is no simple physical explanation for these weak items at present, they seem to represent the collective global evolution of residues, and further research is needed to reveal the significance of these items^29^. Similar findings have been reported in the DCA-based prediction of protein-protein interaction, where the quality of prediction depends on many weak couplings^30^.

It was reported that the motions of orthosteric and allosteric sites are highly correlated^31, 32^, and the regulation between orthosteric and allosteric site is bidirectional^33^. Here we studied the coevolution between orthosteric and allosteric sites and found that their evolutionary coupling strength is stronger than that between orthosteric and other non-functional pockets. We further performed pairwise comparison of the differences in evolutionary coupling scores of residues in the allosteric pocket with orthosteric pockets to identify key allo-residues. Our predicted key allo-residues are supported by previously reported experimental data, such as the predicted key allo-residues in BCR-ABL1, Tar and PDZ3, as well as by cancer mutations. This coevolution-based key allosteric residue prediction method,

KeyAlloSite, could also be used to predict key allosteric residues that can affect the catalytic efficiency of enzymes and are distant from the catalytic site. Taken together, we demonstrated that weak couplings in *J_ij_* contain allosteric information and we developed the first systematic and efficient computational method KeyAlloSite to predict key allo-residues. Our study may contribute to the design and optimization of allosteric drugs, the understanding of allosteric regulatory mechanisms and enzyme engineering.

## Methods and Materials

### The allosteric protein data set

We selected allosteric proteins from the “Core Set” of ASBench^34^ and constructed our data set according to the following criteria: (1) the protein functions as a monomer; (2) the corresponding three-dimensional protein structure data should contain both allosteric ligand and orthosteric ligand; (3) the number of effective homologous sequences of the protein should be greater than 100. Finally, 23 allosteric proteins were selected, including 25 known allosteric sites (Table S1). We collected another two proteins from published literatures, including *E. coli* aspartate chemoreceptor Tar (PDB ID: 4Z9H)^35^ and PDZ3 domain (PDB ID: 1BE9)^36^, as key allo-residues have been reported for these two proteins that can be used for comparative analysis. We used HMMER to search for the homologous sequences of each of the selected allosteric proteins from pfam^37^. Due to the redundancy of homologous sequences, they were reweighted according to the standard of 80% sequence identity to obtain the effective homologous sequences.

### The Evolutionary Coupling Model (ECM)

We used a global statistical model, the evolutionary coupling model (ECM), which can calculate the direct couplings between residues and remove the indirect couplings. The evolutionary coupling model we used here was mainly based on the work of Marks and her co-workers^25, 38^. From the multiple sequence alignment of a protein family, we can calculate the observed frequency 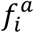 and pairwise co-occurrences 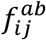 of residues (*a, b*) at position (*i, j*). From this first-order and second-order statistics, we can infer a model to account for the observed statistics optimally, which mainly includes two parameters: single-site propensities *h_i_*(*a*) and direct coevolutionary couplings between residues *J_ij_*(*a,b*). This model defines a probability *P* for each protein sequence ***a*** = (*a*_1_, …, *a_L_*) of length *L*:

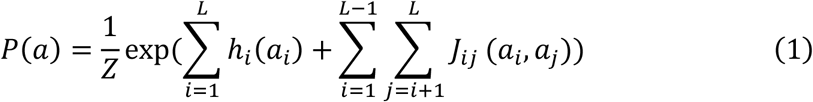

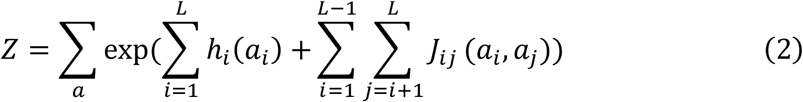

Once the parameters ***h*** and ***J*** are fitted, the Frobenius norm *FN*(*i,j*) of *J_ij_* is used to measure the evolutionary coupling strength between the two sites *i* and *j*.

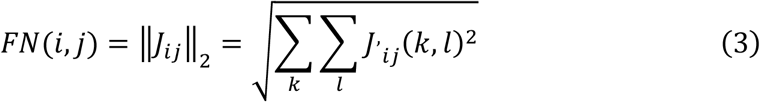

*J’_ij_*. was obtained after *J_ij_* was centralized.

Since the *FN* may include bias caused by phylogeny and under-sampling, it can be corrected with average product correction (APC)^39^. The EC value is the evolutionary coupling score of the two sites.

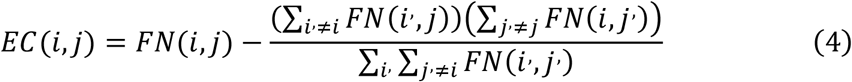

### Evolutionary coupling strength between orthosteric and other pockets

We first performed multiple sequence alignment (Fig. 1A) and calculated the evolutionary couplings between residues using the evolutionary coupling model (Fig. 1B). We then used the CAVITY program ^40, 41^ to identify all the potential ligand binding pockets on the surface of a protein and designated the *m*th pocket as cavity_m. The evolutionary coupling strength (*ECS*_cavity_*m*_) between the orthosteric pocket and the *m*th pocket is defined as the sum of the coupling strength between the residues in the two pockets. The orthosteric pocket here is defined as all of the residues within 6 Å around the orthosteric ligand. If more than 50% of the residues in the *m*th pocket overlap with the orthosteric pocket, the pocket will be excluded. Otherwise, the overlapping residues between the *m*th pocket and orthosteric pocket will be removed from the *m*th pocket, and the remaining residues will be used to calculate the evolutionary coupling strength between the *m*th pocket and orthosteric pocket.

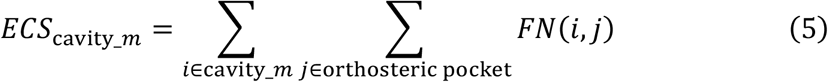

**Fig. 1.**
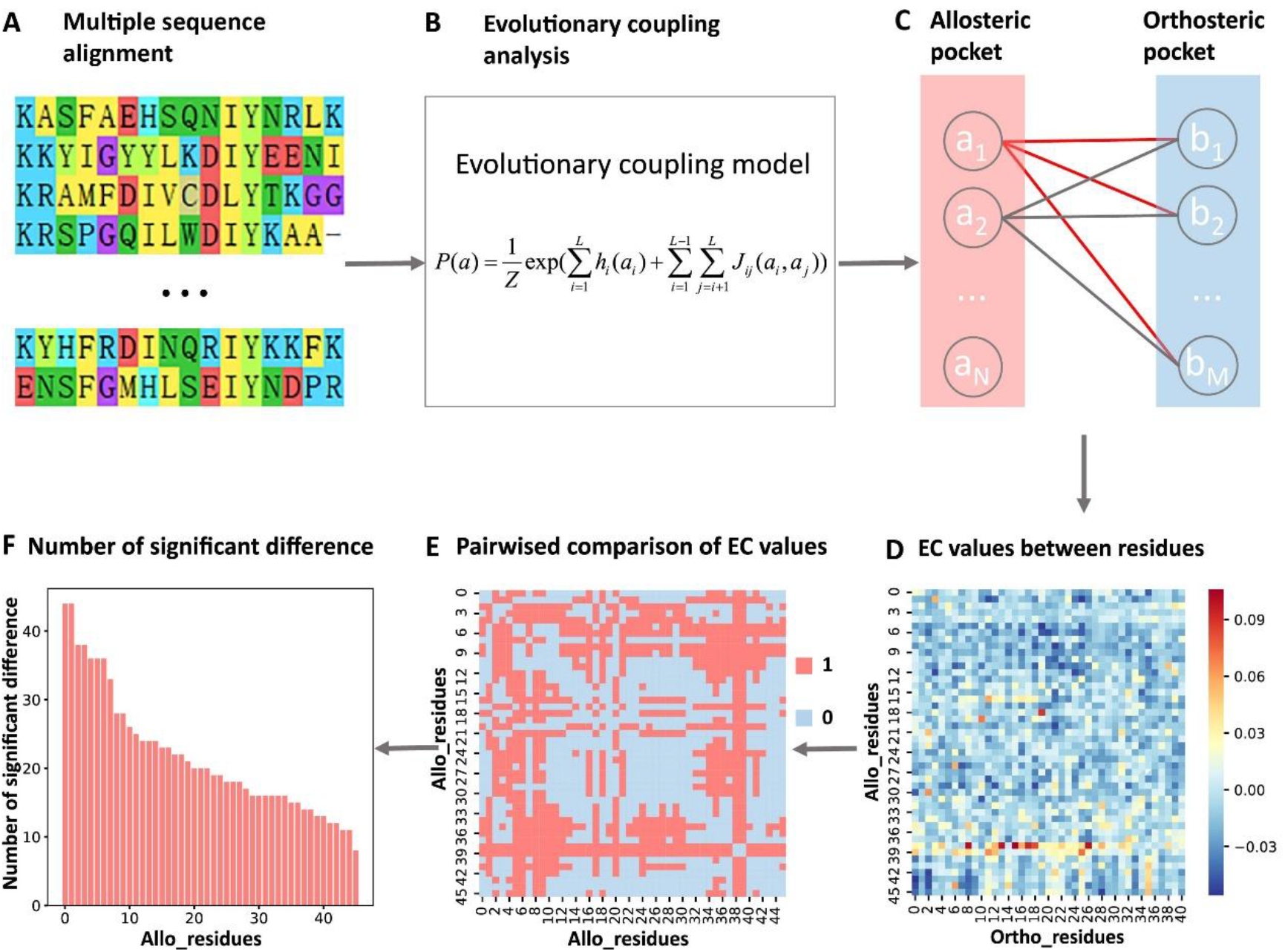
Steps to identify key allo-residues. (A) Multiple sequence alignment. (B) Evolutionary coupling analysis. (C-D) Calculation of the EC values between residues in allosteric and orthosteric pockets. (E) Pairwise compared the difference of EC values corresponding to residues in allosteric pocket. (F) The number of significant differences corresponding to each residue in allosteric pocket.

Finally, we normalized the evolutionary coupling strength of the pockets:

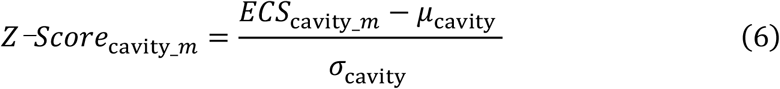

*μ*_cavity_ is the mean value of the evolutionary coupling strength between the orthosteric pocket and the other pockets, and *σ*_cavity_ is the standard deviation of the evolutionary coupling strength.

### Identification of key allo-residues

We first calculated the pairwise EC value between one residue *α_i_* (*i*=1,2,…,*N*) in the allosteric pocket and one residue *b_j_* (*j*=1,2,…*M*) in the orthosteric pocket. An *N* × *M* matrix ***E*** was obtained, and each element *E_ij_* in the matrix ***E*** represents the EC value between residues in the allosteric and orthosteric pockets (Fig. 1C-D). The allosteric pocket here refers to the allosteric pocket found by the CAVITY. In rare cases, if the CAVITY does not find the known allosteric pocket, all of the residues within 8 Å around the allosteric ligand are used as the allosteric pocket. After that, the difference of EC values corresponding to each residue in allosteric pocket was pairwisely compared by using the student’s t-test (α= 0.05), i.e. whether there was a difference between the mean values of each two rows of matrix ***E***. The result of the comparison was expressed by *C_mi_* (*m*=1,2,…,*N*, *i*=1,2,…,*N*), if there was a significant difference, *C_mi_* was assigned a value of 1, if there was no significant difference, *C_mi_* was assigned a value of 0. Finally, an *N* × *N* matrix ***C*** was obtained, and each element in ***C*** indicates whether there was a difference between the residues in the allosteric pocket (Fig. 1E). On this basis, by adding up each column of ***C***, we could get the number of significant differences *d_i_*(*i*=1, 2,…,*N*) between each residue and the remaining residues in the allosteric pocket, i.e. the number of significant differences corresponding to *i*th residue (Fig. 1F).

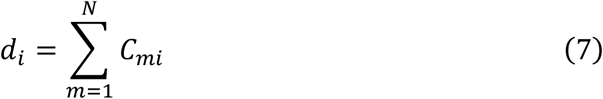

Finally, we normalized the number of significant differences of the residues:

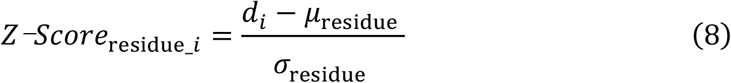

*μ*_residue_ is the mean value of the number of significant differences of the residues, and *σ*_residue_ is the standard deviation of the number of significant differences.

In enzyme catalysis, in addition to residues in the active site, distant residues may also affect enzyme activity, which could not be rationally designed in enzyme engineering. We systematically studied the effect of residues outside of the active site on enzyme catalysis. We calculated the evolutionary coupling value between each residue outside the orthosteric pocket and residues in the orthosteric pocket (i.e. the rows in matrix E are all residues outside the orthosteric pocket here). Using the same steps as above, we determined the number of significant differences *d_i_* for each residue outside the orthosteric pocket and normalized it to Z-score.

## Results

### The evolutionary coupling between orthosteric and allosteric sites is stronger

The sequence lengths of the proteins in our data set range from 166 to 788 amino acid residues (Fig. 2A), and the number of homologous sequences and effective homologous sequences corresponding to each protein were shown in Figure 2B. For each protein in the data set, we used CAVITY to find all the potential ligand binding pockets on the protein surface and explored the evolutionary coupling strength (*ECS*_cavity_m_) between the orthosteric and allosteric pocket, as well as all the other pockets. Among the 25 allosteric pockets in the data set, 23 have Z-Scores greater than 0.5 (Fig. 2C, table S2). The probabilities that the known allosteric pockets were ranked in the top 1, top 2 and top 3 of Z-Scores were 56.0%, 76.0% and 96.0%, respectively (Fig. 2D, table S2), indicating that orthosteric and allosteric pocket are more evolutionarily coupled to each other than the orthosteric and other pockets, which can be used to predict potential allosteric pockets. We further analyzed the two proteins with Z-scores less than 0.5, AR1 and CYP3A4. For AR1, as there were only 108 effective homologous sequences, we speculated that the number of homologous sequences is not enough for evolutionary analysis. The protein sequence based phylogenetic tree of AR1 homologous proteins showed that AR1 located near the tail of the phylogenetic tree (Fig. S1). This implies that this allosteric function did not exist in the early evolutionary period and the allosteric function may have appeared in the later stage of evolution. Due to the relatively large number of sequences in the early evolutionary period and relatively few sequences in the later stage of evolution, the allosteric signal was weak. As a cytochrome P450 protein, CYP3A4 can bind and catalyze the transformation of a variety of substrates. Sequence alignments showed that several positions in its orthosteric pocket are less conserved, which may lead to the difficulty in allosteric site prediction based on evolutional coupling analysis.

**Fig. 2.**
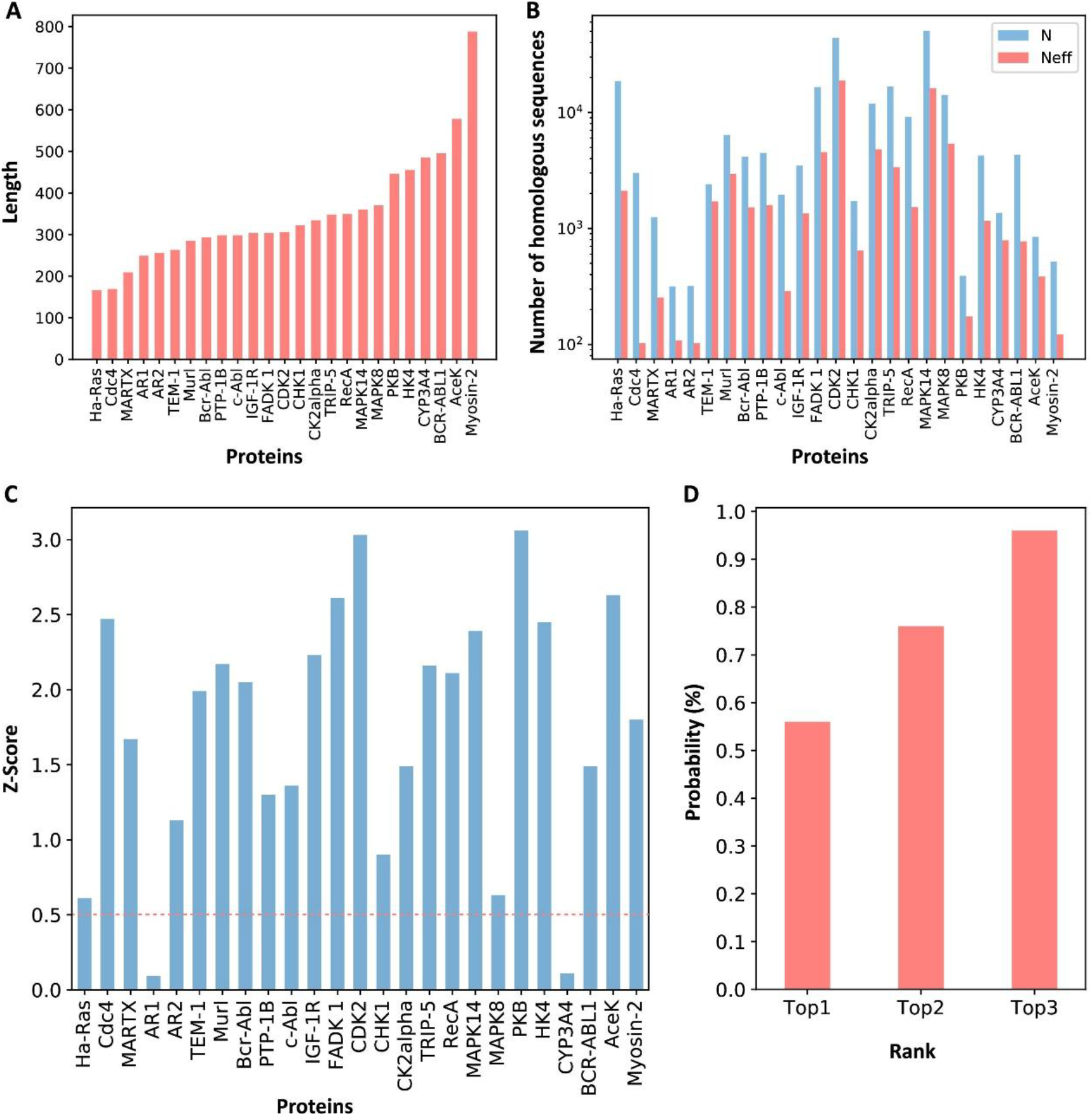
Z-Scores of allosteric pockets and probabilities of ranking an allosteric pocket in the top 3. (A) The sequence lengths of all proteins in our data set. (B) The number of homologous sequences. Neff represents the number of effective homologous sequences obtained under 80% reweighting. (C) Z-Scores of allosteric pockets on proteins in the data set. Among the 25 allosteric pockets, the Z-Scores of 23 allosteric pockets were greater than 0.5. (D) The probabilities that the known allosteric pockets were ranked in the top 1, top 2 and top 3.

We used human Aurora A kinase that is not included in the data set as a test case to further verify whether the evolutionary coupling strength can be used to predict allosteric sites. Aurora A (AurA, PDB ID: 1OL5) is a Ser-Thr protein kinase that is essential for the cell cycle progression. Its abnormal levels can lead to inappropriate centrosome maturation, spindle formation and enhanced cancer growth^42^. AurA is known to be regulated by two distinct allosteric mechanisms, one is specific protein-protein interaction (PPI), which binds TPX2 to its hydrophobic pocket, and the other is phosphorylation of the activation loop at T288 (pT288)^43^. We used CAVITY to find all of the potential ligand binding pockets on the surface of AurA, and a total of 12 pockets were found; cavity_3 is the known allosteric PPI pocket, and cavity_2 is the orthosteric pocket. For consistency, we chose residues within 6 Å around the ATP molecule as the orthosteric pocket. Then we calculated the evolutionary coupling strength between the orthosteric pocket and each of the remaining 11 pockets. Cavity_3 ranked the second among the 11 pockets with a Z-score of 1.48 (Table S3), indicating that the evolutionary coupling strength between the orthosteric and allosteric pockets is indeed stronger. Since phosphorylation mainly occurs on Ser/Thr/Tyr residues, we then calculated the evolutionary coupling strength between the orthosteric pocket and each of the 26 exposed Ser/Thr/Tyr residues and normalized to Z-scores. Among the 26 Ser/Thr/Tyr residues, 10 residues have Z-scores larger than 0.5 and T288 and T287 ranked the 5th and the 4th with a Z-score of 0.83 and 1.05, respectively (Table S3). This indicates that KeyAlloSite can also be used to predict post-translational modification (PTM) sites and the predicted Ser/Thr/Tyr residues with Z-scores greater than 0.5 in addition to T288 and T287 are worth further investigation.

To exclude the possible influence of the pocket size, we further checked the dependence of *ECS*_cavity_m_ on the number of residues used in the calculation. For allosteric and other pockets in each protein, we summed the evolutionary coupling strength of the top 200, top 300 and top 400 residue-residue pairs with the highest *FN*(*i,j*) corresponding to each pocket as the evolutionary coupling strength between each pocket (except for orthosteric pocket) and orthosteric pocket. When different number of residue-residue pairs were used, the Z-Scores corresponding to the evolutionary coupling strength between allosteric and orthosteric pockets in most proteins were still greater than 0.5, which is weakly different from that when all residue-residue pairs were used (Fig. S2). This indicates that the number of residues in the *m*th pocket does not play important role on the evolutionary coupling strength between pockets. In other words, *ECS*_cavity_*m*_ revealed the intrinsic evolutionary coupling between allosteric and orthosteric pockets. These results indicate that the evolutionary coupling strength between orthosteric and allosteric pockets is stronger than that between orthosteric and other pockets, which can be used to predict potential allosteric pockets.

### Coevolution analysis revealed key allo-residues in allosteric pockets

For each protein, we calculated the evolutionary coupling values between the residues in the orthosteric and allosteric pockets by ECM, and compared the corresponding pairwise EC values of the residues in the allosteric pocket. We then calculated the number of significant differences of each residue in the allosteric pocket and normalized to Z-scores. Finally, residues were predicted as key allo-residues if their corresponding Z-scores were greater than 0.8 (Table S4). For the allosteric pockets, the average number of pocket residues is 43, and the average number of identified key allo-residues is 8, accounting for 18.6% of all the residues in the allosteric site (Fig. 3).

**Fig. 3.**
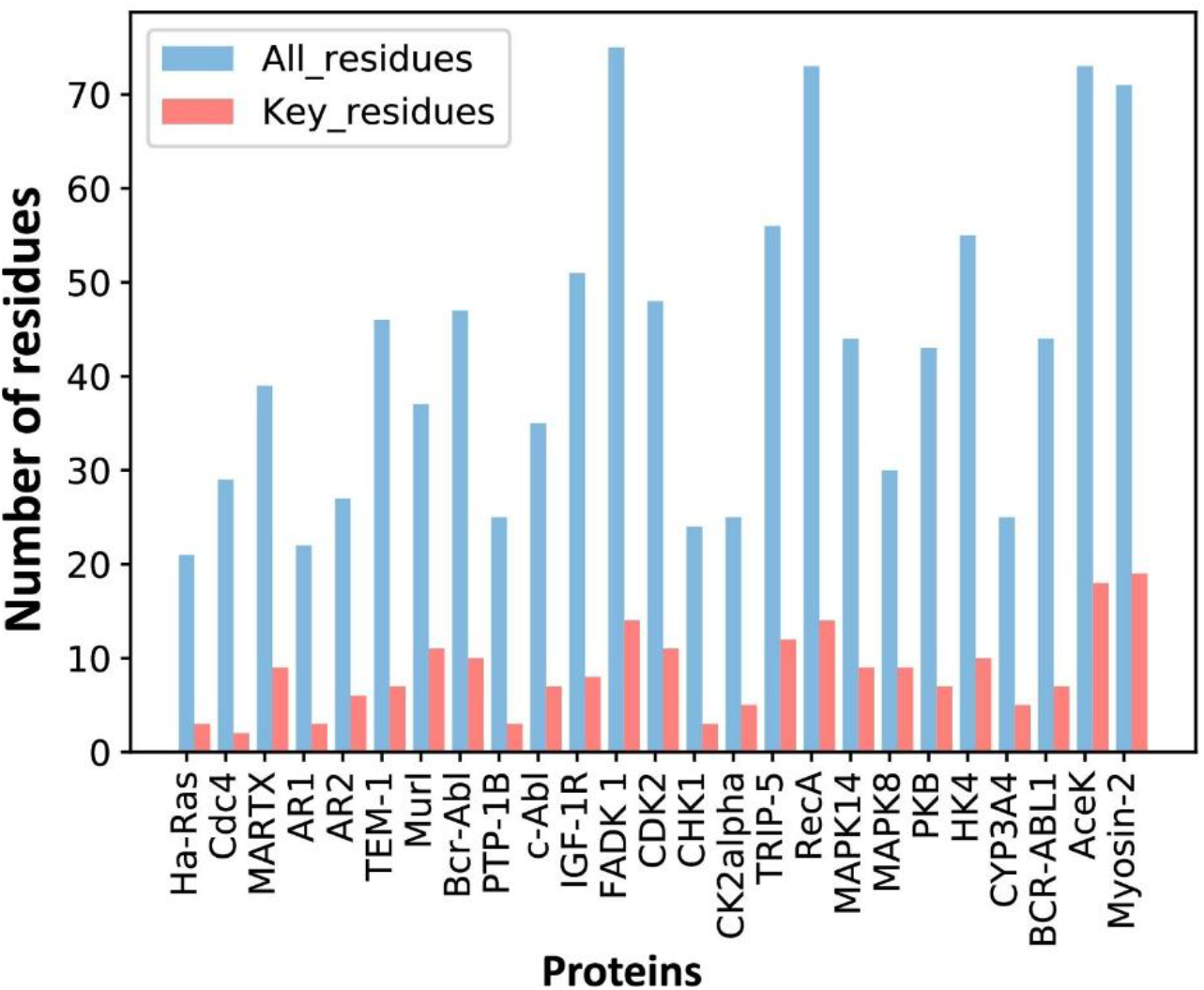
The number of predicted key allo-residues. The number of all residues and predicted key allo-residues in allosteric pockets.

Since the number of homologous sequences is important in coevolution analysis, we selected 7 proteins with a relatively large number of homologous sequences and randomly sampled different numbers of homologous sequences. Since the number of homologous sequences required might be related to the sequence length of the protein itself, the number of homologous sequences was divided by the length of the protein to obtain a ratio. Within this ratio, different ratios of homologous sequences were randomly sampled according to different gradients, and each gradient was repeated 3 times. Then we calculated how many key allo-residues determined by homologous sequences with different gradients were the same as those determined by all homologous sequences. Taking the key allo-residues determined by all homologous sequences as references, we calculated the proportion of the same residues (Fig. S3). It can be seen that generally speaking, 7*L* (± 4*L*) number of effective homologous sequences are enough to give good and stable results. If the protein sequence was relatively short, the number of homologous sequences required could be less; otherwise, more homologous sequences were needed.

### The predicted key allo-residues were supported by experimental results

We searched for literatures to see whether the key allo-residues we predicted were experimentally tested before. The first example is tyrosine-protein kinase ABL1 (BCR-ABL1), which is a fusion protein whose constitutive activity can cause chronic myeloid leukemia (CML). Tyrosine kinase inhibitors targeting the ABL1 ATP-binding site, such as imatinib (Gleevec) and nilotinib (Tasigna), significantly improved the overall survival of CML patients^44, 45^. However, patients may develop drug resistance due to mutations in the ATP-site. The novel fourth generation ABL1 drug, asciminib (ABL001) was developed, which is an allosteric inhibitor that binds to the myristoyl pocket of BCR-ABL1 (Fig. 4A). Asciminib was developed from fragment-based drug discovery approach. In the early stage of hit identification, compounds that bind BCR-ABL1 without inhibition activity were found. Among them, hit **4** binds BCR-ABL1 with a K_d_ of 6 μM. After changing the Cl atom at the *para*-position of the aniline to the CF_3_O-group, hit **5** showed inhibition activity with a slightly weakened K_d_ of 10 μM compared to hit **4** (Fig. 4C)^46^. This shows that the interaction between CF_3_O- and BCR-ABL1 is essential for the allosteric signaling and inhibitory activity. This group forms favorable hydrophobic interaction with L359, one of the key allo-residues predicted by our method (Fig. 4B). In contrast, there is no favorable interaction between hit **4** and L359. These experimental evidences support that the predicted allo-residue L359 plays key role in allosteric signaling.

**Fig. 4.**
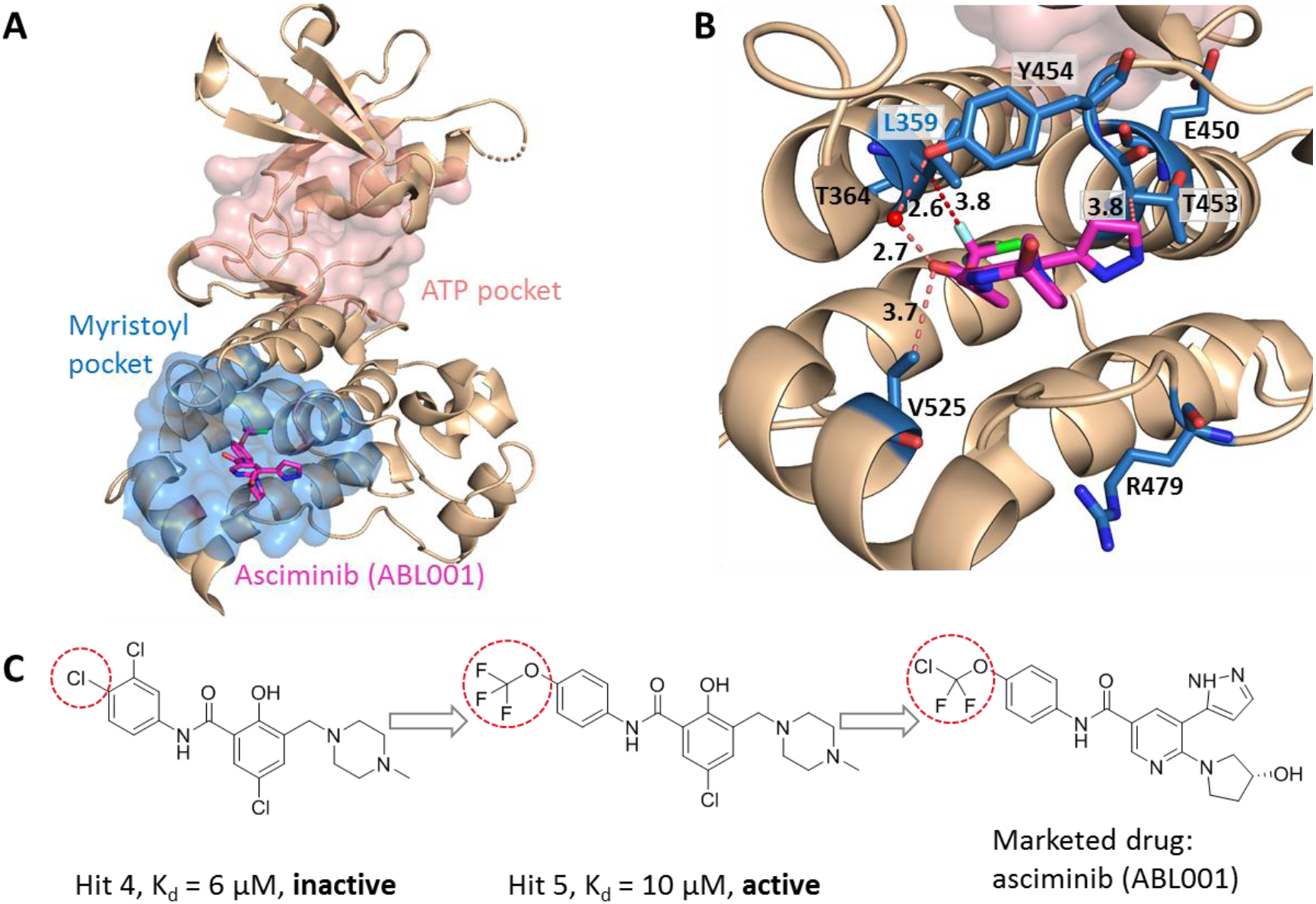
Key allo-residues predicted in BCR-ABL1. (A) The crystal structure of the kinase domain of BCR-ABL1. The allosteric inhibitor asciminib, represented by sticks, binds to the myristoyl pocket (marine). (B) Predicted key allo-residues in the myristoyl pocket. The predicted key allo-residues are represented by marine sticks. One of the predicted key allo-residue L359 forms a favorable hydrophobic interaction with a fluorine atom in asciminib, represented by a red dashed line. Water is represented by a red sphere. (C) The structure of fragment-derived hit 4 and hit 5 and the final marketed drug asciminib.

In the allosteric pocket of the asciminib binding site which contains 44 residues, we predicted 7 key allo-residues. In addition to L359, R479, V525, Y454, E450, T453 and T364 were also identified as potential key allo-residues. T453 forms favorable hydrophobic interaction with the pyrazole ring in asciminib, and Y454 participates in the water-mediated H-bond with the oxygen atom in asciminib. Previous studies have shown that the conformational state of helix-I is important for functional activity and V525 serves as a good indicator for the conformational change. Functional antagonists binding to the myristoyl pocket can bend helix-I and make the disordered region that V525 locates become ordered ^46, 47^. This indicates that V525 plays key role in allosterically regulating ABL1 kinase activity. At the same time, it also shows that coevolutionary information can help to capture multiple allosteric functional conformations of proteins^48^.

### KeyAlloSite correctly identified key allo-residues in other proteins not in the data set

We further tested KeyAlloSite on proteins not included in the data set. The first protein is the *E. coli* aspartate (Asp) chemoreceptor Tar. Tar mediates the chemotaxis of bacteria toward attractants such as Asp, and away from repellents. Tar is a homodimer during transmembrane signaling, and the signals are transmitted from the extracellular region of Tar to the cytoplasmic region through the transmembrane domain^35^. Bi et al. discovered 6 new attractants and 2 new antagonists of Tar by computational virtual screen and experimental study. By comparing the binding patterns of attractants and antagonists, they found that the interactions between the chemoeffectors and Y149 and/or Q152 in Tar are critical for attractant chemotactic signaling^12^. We chose the holo structure that binds Asp (PDB ID: 4Z9H) for analysis, and used the CAVITY to identify all potential ligand binding pockets on the surface of chain B. Among the 3 pockets found, cavity_2 is the pocket where Asp binds, containing 21 residues, which we referred as the allosteric pocket. Since previous studies proposed that transmembrane signaling is triggered by the relative piston-like downward sliding of the α4 helix in the periplasmic domain^35^, we chose the 16 residues (A166-T181) in the C-terminal of the α4 helix as the orthosteric site (Fig. 5A). Through the coevolutionary analysis of residues in the orthosteric and allosteric sites, KeyAlloSite identified 5 key allo-residues in cavity_2, which were I65′, F150′, Y149′, Q152′, P153′ and T154′ (Fig. 5B). It can be seen that our method could predict the key allo-residues Y149 and Q152.

**Fig. 5.**
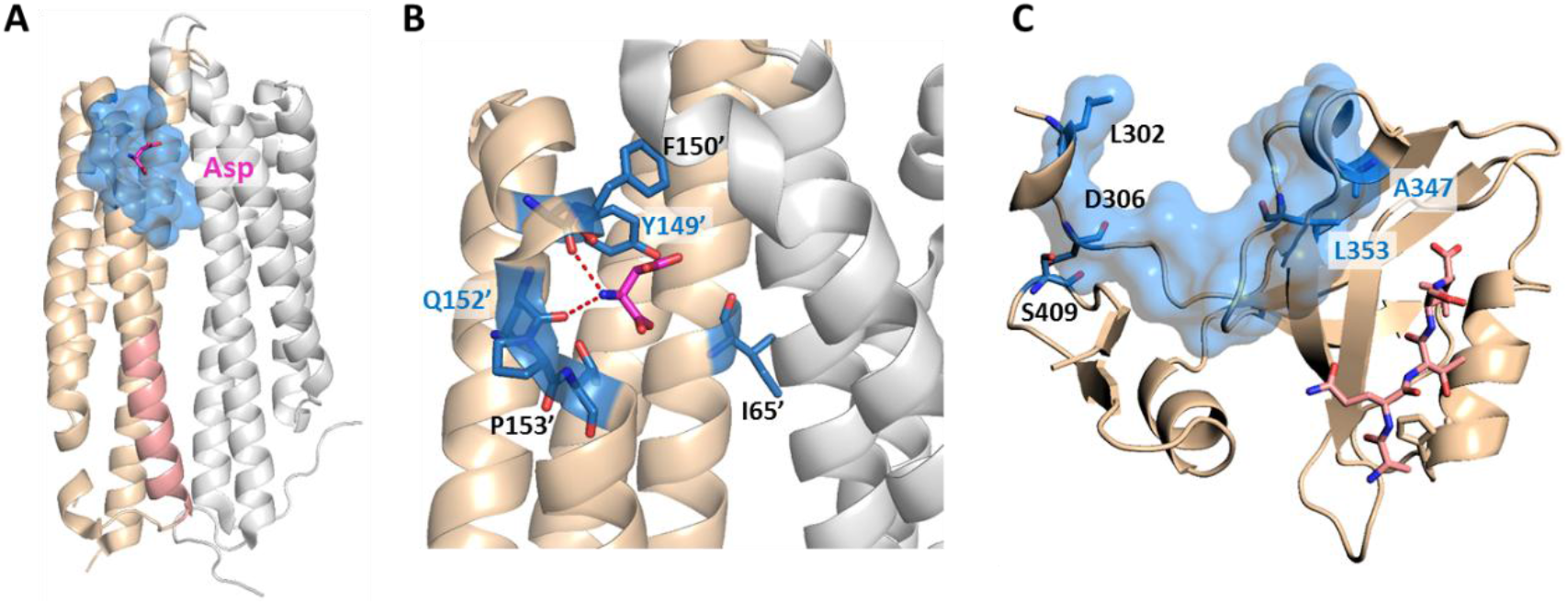
The key allo-residues predicted by our method in Tar and PDZ3. (A) The crystal structure of holo-Tar. Asp is represented by magenta sticks, the allosteric pocket is represented by marine surface, and the salmon helix is selected as the orthosteric site. (B) The key allo-residues predicted at the Asp binding site. The predicted key allo-residues in the allosteric cavity_2 are represented by marine sticks, among which Y149 and Q152 are the true key allo-residues that have been confirmed by experiments. Hydrogen bonds are shown as red dash lines. (C) The predicted key allo-residues in PDZ3. The peptide bound to the orthosteric site is represented by salmon sticks, the allosteric pocket is represented by marine surface, and the predicted key allo-residues are represented by marine sticks.

The second protein is the PDZ3 domain, and its allosteric mechanism has been extensively studied. We selected the crystal structure of PDZ3 binding with a peptide in its orthosteric site for analysis (PDB ID: 1BE9)^36^. Since the allosteric site of this structure does not bind an allosteric ligand, we used CAVITY to find the potential ligand binding pockets on its surface. Among the three pockets identified, cavity_1 is the orthosteric pocket and cavity_2 contains the known allosteric sites, which were used for further analysis. KeyAlloSite predicted D306, S409, L302, A347 and L353 as key allo-residues (Fig. 5C). Kalescky et al. used rigid residue scan to identify residues that are important for the allosteric effect of the PDZ3 domain. In the rigid residue scan, only one residue was regarded as a single rigid body in each molecular dynamics simulation. They proposed that A347 is a “switch residue”, which is needed to turn on the allosteric effect^20^. Lockless et al. used evolutionary data of protein families to measure the statistical coupling between amino acid positions. For the PDZ protein family, they found that there is strong statistical couplings between A347 and L353 and the key residue H372 of the orthosteric site, and verified using thermodynamic mutational studies^49^. Moreover, McLaughlin et al. developed a high-throughput quantitative method that can individually replace a residue at each position with every other residue for comprehensive single-mutation studies. Their results showed that mutations of A347 and L353 caused significant functional loss^50^. These evidences all indicate that the key allo-residues A347 and L353 we predicted are important for the protein function by allosteric regulation.

### KeyAlloSite identified pathogenetic mutations in human proteins

Previous studies have shown that allosteric mutation, that is, abnormal protein allosteric regulation caused by mutation, is related to pathological processes such as cancer^51^. Shen et al. analyzed the dysfunction of allosteric proteins caused by somatic mutations in about 7000 cancer genomes across 33 cancer types, mapped these mutations to allosteric sites, orthosteric sites and other sites in the Allosteric Database, and established the Allo-Mutation dataset^52^. We searched for the somatic mutation data corresponding to the human proteins in our data set from the Allo-Mutation dataset, and found that some of the predicted key allo-residues in 7 of the 15 human proteins were mutated in a variety of cancers (Table 1). Among them, cancers that contain a large number of mutations in key allo-residues are uterine corpus endometrial carcinoma (UCEC) and skin cutaneous melanoma (SKCM). This indicates that the abnormal allosteric regulation caused by the mutation of key allo-residues plays key role in the occurrence and development of cancer. These key allo-residues can affect allosteric signal transduction and thus affect protein functions, suggesting that KeyAlloSite can be used to predict key pathogenetic mutations in proteins.

**Table 1.** Predicted key allo-residues that were mutated in cancers.

| Protein name | Gene name | Predicted key allo-residues | Mutation<sup>a</sup> | Cancer type<sup>b</sup> |
| --- | --- | --- | --- | --- |
| AR1 | AR | D732 | D732N | SKCM |
| AR2 | AR | M832 | M832I | SKCM |
| PTP-1B | PTPN1 | M282 | M282T | COAD |
| CDK2 | CDK2 | P155 | P155H | UCEC |
| CK2alpha | CSNK2A1 | F54; A110 | F54C; A110T | UCEC; UCEC, GBM |
| MAPK14 | MAPK14 | P191; E192 | P191S; P191H; E192Q | SKCM; KIRC; BLCA |
| MAPK8 | MAPK8 | E195; M200 | E195K; M200I | UCEC; SKCM |
| CYP3A4 | CYP3A4 | F219 | F219L | UCEC |

<sup>a</sup>Mutation: confirmed disease mutations among the predicted key allo-residues; <sup>b</sup>Cancer type: COAD: Colon adenocarcinoma; SKCM: Skin cutaneous melanoma; UCEC: Uterine corpus endometrial carcinoma; GBM: Glioblastoma multiforme; KIRC: Kidney renal clear cell carcinoma; BLCA: Bladder urothelial carcinoma.

### KeyAlloSite can also identify key allosteric functional residues of enzymes

Enzyme evolution studies mainly focus on mutations in the substrate binding pocket ^53,54^. However, covariant residues far from the substrate binding site may also play important role in regulating catalysis, which are difficult to identify. Since KeyAlloSite can calculate the functional coupling between residues outside and inside the orthosteric pocket, we wonder whether it can also be used to identify key allo-residues for enzyme activity regulation. We used *Candida antarctica* lipase B (CALB, PDB ID: 1TCA), which has many annotated key functional residues^55^, as a test case. CALB is one of the most widely used biocatalysts in academia and industry that is often applied in acylating kinetic resolution of racemic alcohols and amines and desymmetrization of diols and diacetates, and is robust and easy to express^54^. We selected the 6 angstrom-cutoff orthosteric pocket, and calculated the EC values between each residue outside the orthosteric pocket and the residues in the orthosteric pocket. Then we compared the difference in EC values corresponding to each residue outside the orthosteric pocket, and calculated the significant difference number *d_i_* corresponding to each residue. Finally, the significant difference number *d_i_* was normalized to Z-score. Due to the large number of residues outside the orthosteric pocket, residues with Z-scores greater than 0.9 were referred as key allo-residues here. Our method predicted a total of 52 residues from the 296 residues outside the orthosteric pocket, of which 20 residues have been annotated as functional residues in literature according to mutagenesis experiments (Fig. 6A, Table S5). For example, A225 ranked the third out of the 52 predicted residues with a Z-Score of 3.04, and A225M improves the catalytic efficiency of the enzyme by about 11 folds^55^. Therefore, our method can also be used to predict allosteric functional residues that are important for enzyme catalysis, providing a new computational tool for identifying mutant enzyme with improved catalytic properties.

**Fig. 6.**
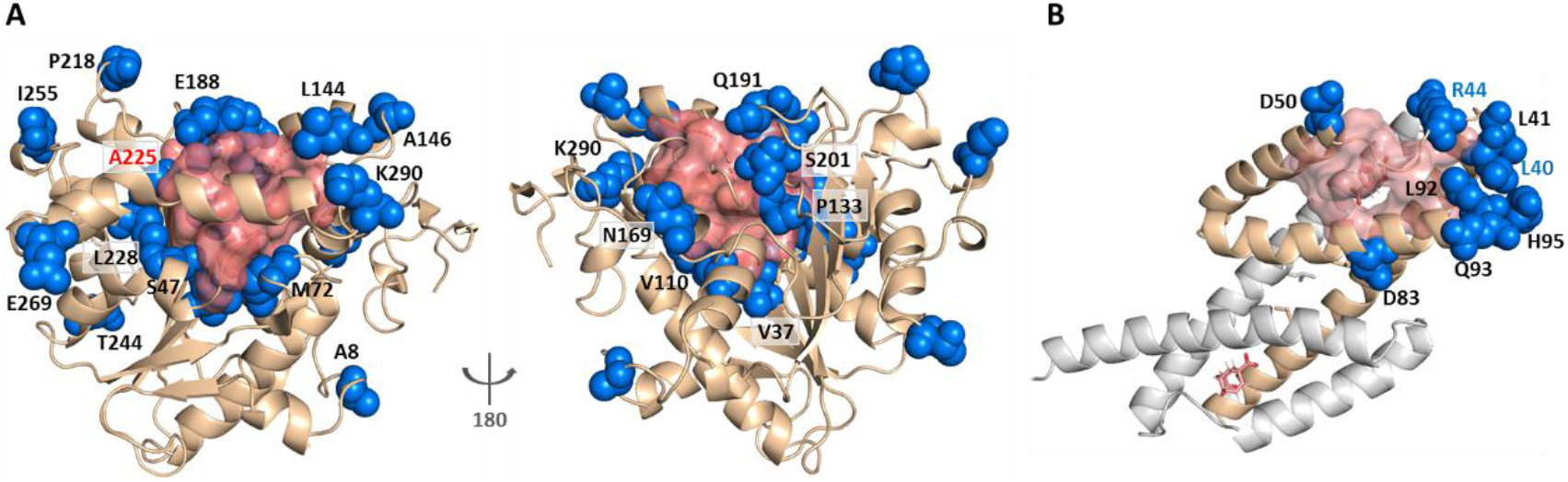
KeyAlloSite predicted key allo-residues for enzymes. (A) KeyAlloSite predicted key allo-residues for *Candida antarctica* lipase B. Among the predicted residues, the residues that have been annotated by the literature are shown as marine spheres, and the orthosteric pocket is represented by salmon surface. (B) KeyAlloSite predicted key allo-residues for *E. coli* chorismate mutase. Experimentally discovered key functional residues of CM are shown as marine spheres, the labels of key allo-residues predicted by KeyAlloSite are shown in marine, and the orthosteric pocket and ligand are represented by salmon surface and sticks.

Russ *et al*. recently used an evolution-based model to design chorismate mutase enzymes (CMs). They used DCA to learn the constraints for specifying proteins purely from evolutionary sequence data, and performed Monte Carlo sampling from this model to generate artificial sequences. They were able to obtain proteins with natural-like catalytic function with sequence diversity. Eight residues (L40, L41, R44, D50, D83, L92, Q93, H95) at the periphery of the active site were found to be important for CM catalysis in *E. coli* specific function^29^ (Fig. 6B). We wonder if some of these residues regulate the enzyme function by allostery. We used the 6 angstrom-cutoff orthosteric pocket and identified 46 residues on the surface of the protein outside the orthosteric pocket. We calculated the EC values between each of these 46 residues and the orthosteric pocket and compared the difference in EC values of these 46 residues. We then calculated the significant difference number *d_i_* corresponding to each of these 46 residues and normalized to Z-scores. Using the criterion of Z-score > 0.9, we predicted 10 key allo-residues from the 48 residues, which were L86, A9, R44, Q89, E57, E12, L40, D69, R58, and N5. Among them, two residues (R44 and L40) were shown to be important for CM catalysis by Russ *et al*. Russ *et al*. used DCA to show that sequence-based statistical models contain protein function information, while our study used DCA to predict allosteric sites or key allosteric residues based on the significance analysis of evolutionary coupling between these sites with the orthosteric site. Their study showed that the function of proteins seems to depend on many weak terms in *J_ij_*, which lack simple physical interpretations. Our study demonstrated that weak terms in *J_ij_* contain information for protein allosteric function evolution and can be used to predict allosteric sites and key allosteric residues.

## Discussion

Identifying key allosteric residues responsible for allosteric signaling is important for the design and optimization of allosteric drugs, enzyme and protein engineering studies. We developed, KeyAlloSite, a novel method for predicting allosteric sites and key allo-residues based on the evolutionary coupling model. To the best of our knowledge, this is the first systematic and efficient computational method to predict key allo-residues. Our study demonstrated that orthosteric and allosteric pockets are coupled in protein function evolution. Our predicted key allo-residues are in accordance with previously reported experimental studies in the BCR-ABL1, Tar and PDZ3 systems, as well as key cancer mutations. We further showed that KeyAlloSite can be applied to predict key allosteric residues distant from the catalytic site that are important for enzyme catalysis. Our study also gives a possible physical explanation for the weak couplings in *J_ij_*, that is, they may represent allosteric functional couplings. The predicted key allo-residues can help us to understand the mechanism of allosteric regulation, to provide reference and guidance for the rational design and optimization of allosteric drugs and to facilitate enzyme engineering.

It should be noted that although we used the three-dimensional structure of proteins and their binding ligands in our analysis, KeyAlloSite can also be applied in cases where no three-dimensional structures are available on condition that a certain number of homologous sequences of the protein under investigation and location of the functional site are known. Along with the rapid progress in recent years, protein structure prediction methods such as AlphaFold^56^ can be used to predict the protein structure first. At the same time, as our method calculates the evolutionary coupling between any residue of the protein and residues of the orthosteric pocket, KeyAlloSite can be used to predict not only key allosteric residues, but also post-translational modification sites that have functional correlation with orthosteric sites, which will be further studied in the future.

## Data availability

The data that support the results of this study are in the Supplementary Data including information of the allosteric proteins in the data set (Table S1); list of the Z-Scores and ranking of allosteric pockets in the data set (Table S2); KeyAlloSite prediction results of Aurora A kinase (Table S3); list of the predicted key allo-residues in allosteric pockets (Table S4); the key allo-residues predicted by our method on *Candida antarctica* lipase B (Table S5); phylogenetic tree of androgen receptor (Figure S1); comparison of evolutionary coupling strength between pockets when all residue pairs and partial residue pairs were used (Figure S2); and random sampling of homologous sequences (Figure S3).

## Acknowledgements

The authors thank Jintao Zhu from the Center for Quantitative Biology and Gaoxiang Pan from the College of Chemistry and Molecular Engineering, Peking University for helpful discussions. This study was supported in part by the Chinese Academy of Medical Sciences (2021-I2M-5-014) and National Natural Science Foundation of China (21633001).

## Author contributions

Juan Xie, Investigation, Conceptualization, Data curation, Methodology, Formal analysis, Visualization, Writing – original draft preparation, Writing – review & editing; Weilin Zhang, Conceptualization, Methodology, Investigation, Formal analysis; Xiaolei Zhu, Methodology, Writing – review & editing; Minghua Deng, Conceptualization, Formal analysis, Methodology; Luhua Lai, Conceptualization, Methodology, Writing – review & editing, Supervision, Funding acquisition.

**Correspondence** and requests for materials should be addressed to Prof. Luhua Lai.

