## Supplementary Information for "Coevolution-based prediction of key allosteric residues for protein function regulation"

[1] Center for Quantitative Biology, Academy for Advanced Interdisciplinary Studies, Peking University, Beijing, China. [2] BNLMS, Peking-Tsinghua Center for Life Sciences at the College of Chemistry and Molecular Engineering, Peking University, Beijing, China. [3] School of Sciences, Anhui Agricultural University, Hefei, Anhui, China. [4] School of Mathematical Sciences, Peking University, Beijing, China. [5] Center for Statistical Science, Peking University, Beijing, China. [6] Research Unit of Drug Design Method, Chinese Academy of Medical Sciences (2021RU014), Beijing, China.

**Supplementary material:**

**Table supplement 1**. Information of the allosteric proteins in the data set

**Table supplement 2**. List of the Z-scores and ranking of allosteric pockets in the data set

**Table supplement 3**. KeyAlloSite prediction results of Aurora A kinase

**Table supplement 4**. List of the predicted key allo-residues in allosteric pockets

**Table supplement 5**. The key allo-residues predicted by our method on *Candida antarctica* lipase B

**Figure supplement 1**. Phylogenetic tree of the androgen receptor.

**Figure supplement 2**. Comparison of evolutionary coupling strength between pockets when all residue pairs and partial residue pairs were used.

**Figure supplement 3**. Random sampling of homologous sequences.

**Table supplement 1**. Information of the allosteric proteins in the data set

| Protein name | PDB ID_AS[a] | PDB ID_OS | Allosteric effector | Orthosteric effector | Organism | Allosteric type |
|---|---|---|---|---|---|---|
| Androgen receptor (AR1) | 2YHD | 2PIO | AV6 | DHT | Homo sapiens | Inhibitor |
| Androgen receptor (AR2) | 2YLO | 2PIO | YLO | DHT | Homo sapiens | Inhibitor |
| Beta-lactamase TEM (TEM-1) | 1PZO | 1AXB | CBT | FOS | Escherichia coli | Inhibitor |
| Casein kinase II subunit alpha (CK2alpha) | 3H30 | 3H30 | RFZ | RFZ | Homo sapiens | Inhibitor |
| Cell division control protein 4 (Cdc4) | 3MKS | 1NEX | C1C | TPO | Saccharomyces cerevisiae | Inhibitor |
| Cyclin-dependent kinase 2 (CDK2) | 3PXF | 1B38 | 2AN | ATP | Homo sapiens | Inhibitor |
| Cytochrome P450 3A4 (CYP3A4) | 1W0F | 1W0F | STR | HEM | Homo sapiens | Regulator |
| Focal adhesion kinase 1 (FADK 1) | 4EBW | 2IJM | 0PF | ATP | Homo sapiens | Inhibitor |
| Glucokinase (HK4) | 1V4S | 3F9M | MRK | GLC | Homo sapiens | Activator |
| Glutamate racemase (MurI) | 2JFN | 2JFZ | UMA | DGL | Escherichia coli | Activator |
| Gtpase hras (Ha-Ras) | 4DLR | 4DLR | DTU | GNP | Homo sapiens | Regulator |
| Insulin-like growth factor 1 receptor (IGF-1R) | 3LW0 | 1JQH | CCX | ANP | Homo sapiens | Inhibitor |
| Isocitrate dehydrogenase kinase/phosphatase (AceK) | 3LCB | 3LCB | AMP | ATP | Escherichia coli | Inhibitor |
| Kinesin-like protein kif11 (TRIP-5) | 3ZCW | 3ZCW | 4A2 | ADP | Homo sapiens | Inhibitor |
| Mitogen-activated protein kinase 14 (MAPK14) | 4E6C | 3KF7 | 0O8 | L9G | Homo sapiens | Activator |
| Mitogen-activated protein kinase 8 (MAPK8) | 3O2M | 1UKI | 46A | 537 | Homo sapiens | Inhibitor |
| Myosin-2 heavy chain (Myosin-2) | 2JHR | 1YV3 | PBQ | ADP | Dictyostelium discoideum | Inhibitor |
| Protein RecA (RecA) | 2G88 | 2G88 | DTP | DTP | Mycolicibacterium smegmatis | Regulator |

Table S1. Continued

| | | | | | | |
|---|---|---|---|---|---|---|
| Protein-tyrosine phosphatase 1B (PTP-1B) | 1T49 | 1BZC | 892 | TPI | Homo sapiens | Inhibitor |
| RAC-alpha serine/threonine-protein kinase (PKB) | 3O96 | 4EKK | IQO | ANP | Homo sapiens | Inhibitor |
| RTX toxin RtxA (MARTX) | 3GCD | 3GCD | IHP | AZ0 | Vibrio cholerae | Regulator |
| Serine/threonine-protein kinase Chk1 (CHK1) | 3F9N | 2E9N | 38M | 76A | Homo sapiens | Inhibitor |
| Tyrosine protein kinase ABL1 (Bcr-Abl) | 3K5V | 3K5V | STJ | STI | Mus musculus | Inhibitor |
| Tyrosine-protein kinase ABL1 (BCR-ABL1) | 5MO4 | 5MO4 | AY7 | NIL | Homo sapiens | Inhibitor |
| Tyrosine-protein kinase ABL1 (c-Abl) | 3PYY | 3PYY | 3YY | STI | Homo sapiens | Activator |

[a]AS, allosteric site; OS, orthosteric site; PDB ID_AS, protein structure in the allosteric effector-bound state; PDB ID_OS, protein structure used to define OS.

**Table supplement 2**. List of the Z-scores and ranking of allosteric pockets in the data set

| Protein name | Z-score[a] | Rank[b] |
|---|---|---|
| Cdc4 | 2.47 | 1/11 |
| MARTX | 1.67 | 1/4 |
| Ha-Ras | 0.61 | 2/3 |
| TEM-1 | 1.99 | 1/7 |
| MurI | 2.17 | 1/7 |
| AR1 | 0.09 | 3/7 |
| IGF-1R | 2.23 | 1/8 |
| Bcr-Abl | 2.05 | 1/10 |
| PTP-1B | 1.30 | 3/12 |
| c-Abl | 1.36 | 3/11 |
| AR2 | 1.13 | 2/7 |
| FADK 1 | 2.61 | 1/8 |
| CDK2 | 3.03 | 1/15 |
| CHK1 | 0.90 | 3/9 |
| CK2alpha | 1.49 | 1/15 |
| TRIP-5 | 2.16 | 1/11 |
| RecA | 2.11 | 2/13 |
| MAPK14 | 2.39 | 1/11 |
| MAPK8 | 0.63 | 3/16 |
| PKB | 3.06 | 1/12 |
| HK4 | 2.45 | 1/10 |
| CYP3A4 | 0.11 | 8/14 |
| BCR-ABL1 | 1.49 | 2/15 |
| AceK | 2.63 | 1/18 |
| Myosin-2 | 1.80 | 2/24 |

[a]Z-score: the Z-score corresponding to the evolutionary coupling strength of the allosteric pocket. [b]Rank: the ranking of allosteric pocket in all pockets except orthosteric pockets according to Z-score in KeyAlloSite.

**Table supplement 3**. KeyAlloSite prediction results of Aurora A kinase

| PPI | | Phosphorylation | | |
|---|---|---|---|---|
| Pockets | Z-score | Residues | Sites | Z-score |
| cavity_1 | 1.95 | THR | 235 | 1.36 |
| **cavity_3** | **1.48** | SER | 245 | 1.23 |
| cavity_6 | 0.80 | SER | 249 | 1.09 |
| cavity_7 | 0.50 | **THR** | **287** | **1.05** |
| cavity_9 | -0.27 | **THR** | **288** | **0.83** |
| cavity_12 | -0.39 | THR | 204 | 0.82 |
| cavity_11 | -0.51 | SER | 361 | 0.70 |
| cavity_10 | -0.54 | TYR | 148 | 0.67 |
| cavity_8 | -0.56 | SER | 369 | 0.60 |
| cavity_4 | -1.11 | SER | 226 | 0.60 |
| cavity_5 | -1.34 | SER | 155 | 0.49 |
| | | TYR | 212 | 0.47 |
| | | TYR | 219 | 0.32 |
| | | THR | 384 | 0.17 |
| | | THR | 353 | 0.12 |
| | | THR | 347 | 0.06 |
| | | TYR | 338 | -0.26 |
| | | TYR | 334 | -0.29 |
| | | SER | 283 | -0.34 |
| | | THR | 292 | -0.44 |
| | | SER | 284 | -0.48 |
| | | SER | 266 | -1.33 |
| | | THR | 333 | -1.33 |
| | | SER | 342 | -1.91 |
| | | SER | 387 | -1.97 |
| | | SER | 123 | -2.23 |

**Table supplement 4**. List of the predicted key allo-residues in allosteric pockets

| Protein | Predicted key allo_residues[a] |
|---|---|
| Ha-Ras | A/62,A/89,A/94, |
| Cdc4 | D/662,D/634, |
| MARTX | A/200,A/135,A/85,A/54,A/51,A/201,A/168,A/180,A/167, |
| AR1 | A/736,A/732,A/725, |
| AR2 | A/724,A/833,A/722,A/832,A/721,A/826, |
| TEM-1 | A/246,A/245,A/217,A/214,A/248,A/233,A/216, |
| MurI | A/229,A/237,A/233,A/150,A/123,A/96,A/226,A/151,A/116,A/235,A/101, |
| Bcr-Abl | A/523,A/453,A/450,A/352,A/356,A/479,A/360,A359,A358,A351 |
| PTP-1B | A/195,A/187,A/282, |
| c-Abl | A/478,A/359,A/519,A/482,A/455,A/364,A/481, |
| IGF-1R | A/1136,A/1135,A/1167,A/1154,A/1039,A/1155,A/1045,A/1156, |
| FADK 1 | A/546,A/621,A/618,A/540,A/598,A/475,A/545,A/535,A/474,A/544,A/541,A/668,A/472,A/548, |
| CDK2 | A/147,A/51,A/154,A/125,A/148,A/158,A/157,A/35,A/155,A/71,A/61, |
| CHK1 | A/233,A/132,A/205, |
| CK2alpha | A/54,A/104,A/59,A/36,A/110, |
| TRIP-5 | A/115,A/117,A/134,A/235,A/232,A/202,A/162,A/227,A/213,A/164,A/116,A/170, |
| RecA | A/54,A/328,A/50,A/52,A/337,A/257,A/256,A/247,A/255,A/56,A/254,A/49,A/246,A/53, |
| MAPK14 | A/200,A/192,A/191,A/296,A/194,A/240,A/293,A/207,A/193, |
| MAPK8 | A/197,A/255,A/257,A/196,A/185,A/251,A/195,A/192,A/200, |
| PKB | A/273,A/331,A/202,A/275,A/213,A/18,A/14, |
| HK4 | A/457,A/460,A/69,A/461,A/459,A/96,A/453,A/248,A/207,A/201, |
| CYP3A4 | A/484,A/482,A/308,A/304,A/219, |
| BCR-ABL1 | A/479,A/525,A/454,A/450,A/359,A/453,A/364, |
| AceK | A/59,A/56,A/291,A/256,A/414,A/412,A/372,A/294,A/371,A/299,A/114,A/104,A/374,A/361,A/298,A/384,A/120,A/100, |
| Myosin-2 | A/240,A/468,A/267,A/264,A/471,A/435,A/262,A/259,A/239,A/635,A/430,A/632,A/594,A/588,A/263,A/261,A/587,A/426,A/438, |

[a]Predicted key allo_residues: Chain/Residue sequence number

**Table supplement 5**. The key allo-residues predicted by our method on *Candida antarctica* lipase B

| Key allo-residues[a] | Z-score | Key allo-residues[a] | Z-score |
|---|---|---|---|
| **P133** | 3.98 | A305 | 1.58 |
| L73 | 3.94 | G44 | 1.54 |
| **A225** | 3.04 | **A146** | 1.52 |
| **S201** | 3.04 | G60 | 1.52 |
| S150 | 3.04 | T186 | 1.46 |
| **E188** | 3.00 | R249 | 1.44 |
| **E269** | 2.72 | **T244** | 1.44 |
| **S47** | 2.51 | S153 | 1.44 |
| **N169** | 2.33 | **V37** | 1.42 |
| S250 | 2.31 | R309 | 1.36 |
| V78 | 2.31 | W113 | 1.34 |
| **M72** | 2.31 | S195 | 1.30 |
| P152 | 2.25 | L90 | 1.21 |
| S67 | 2.25 | F205 | 1.19 |
| S243 | 2.21 | L204 | 1.15 |
| L140 | 2.13 | P63 | 1.09 |
| T238 | 2.05 | Q112 | 1.01 |
| **V110** | 2.05 | **I255** | 0.99 |
| **Q191** | 1.99 | L167 | 0.99 |
| **K290** | 1.95 | **L144** | 0.97 |
| **L228** | 1.93 | L128 | 0.97 |
| **P218** | 1.88 | T88 | 0.97 |
| S184 | 1.86 | **A8** | 0.97 |
| I66 | 1.82 | Y203 | 0.95 |
| P260 | 1.80 | S161 | 0.93 |
| **D187** | 1.76 | I87 | 0.93 |

[a]Key allo-residues: Among the predicted key allo-residues, the residues that have been annotated by the literature are marked in bold.

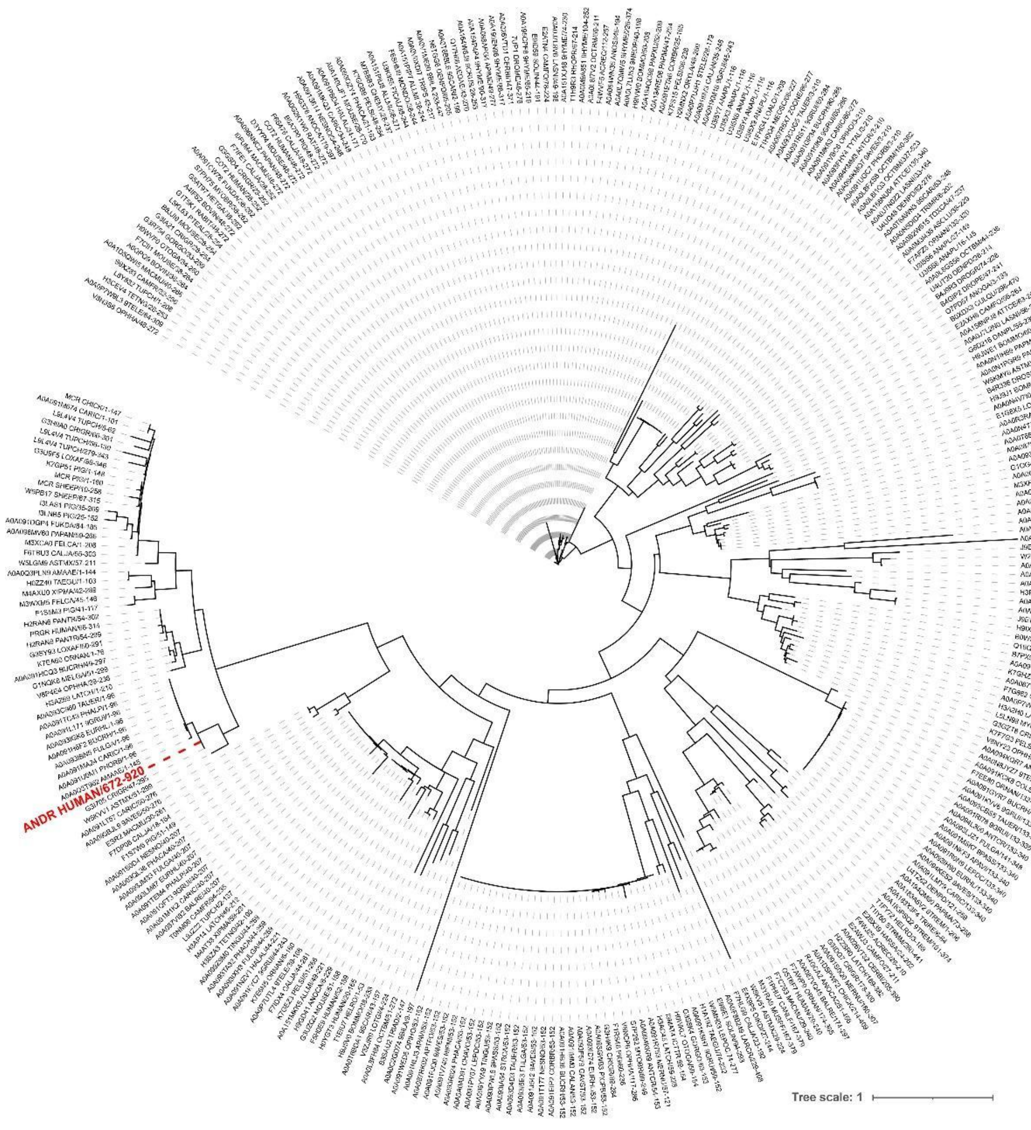


**Figure supplement 1**. Phylogenetic tree of the androgen receptor.

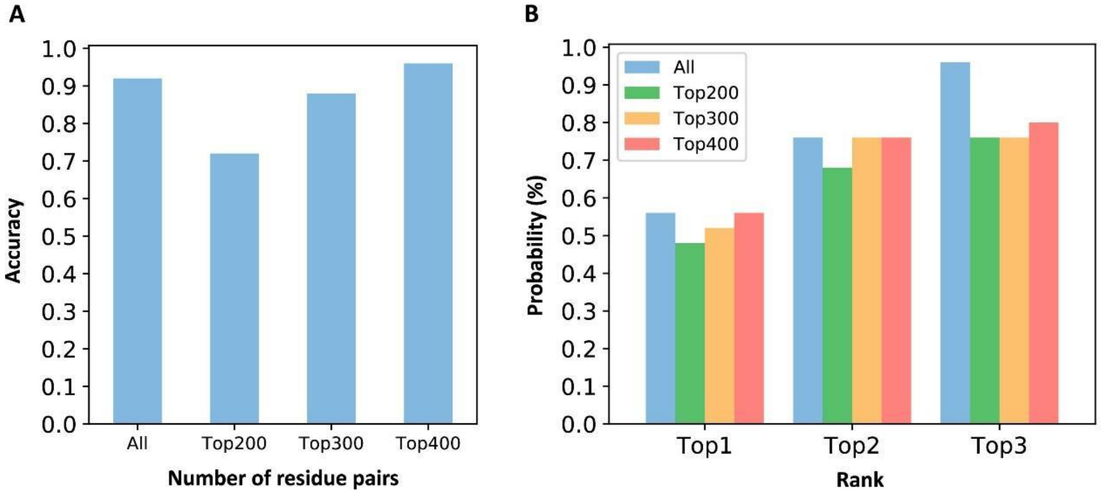


**Figure supplement 2**. Comparison of evolutionary coupling strength between pockets when all residue pairs and partial residue pairs were used. To further check the influence of the number of residues in the $m$th pocket on $ECS_{\text{cavity\_}m}$, for allosteric and other pockets on each protein, we summed the evolutionary coupling strength of top 200, top 300 and top 400 residue-residue pairs with the highest FN$(i, j)$ corresponding to each pocket as the evolutionary coupling strength between each pocket and orthosteric pocket. (A) Prediction accuracy of using different numbers of residue pairs. We defined that the criterion for successful prediction is that the Z-score of the allosteric pocket is greater than 0.5. It can be seen that there is a small difference between the prediction accuracy when using all residue pairs and using different number of residue pairs. (B) The probability that the allosteric pocket is ranked in the top 3 using different numbers of residue pairs. It can be seen that there is a small difference between the probability that the allosteric pocket was ranked in the top 3 when all residue pairs and different number of residue pairs were used, especially in top 1 and top 2.

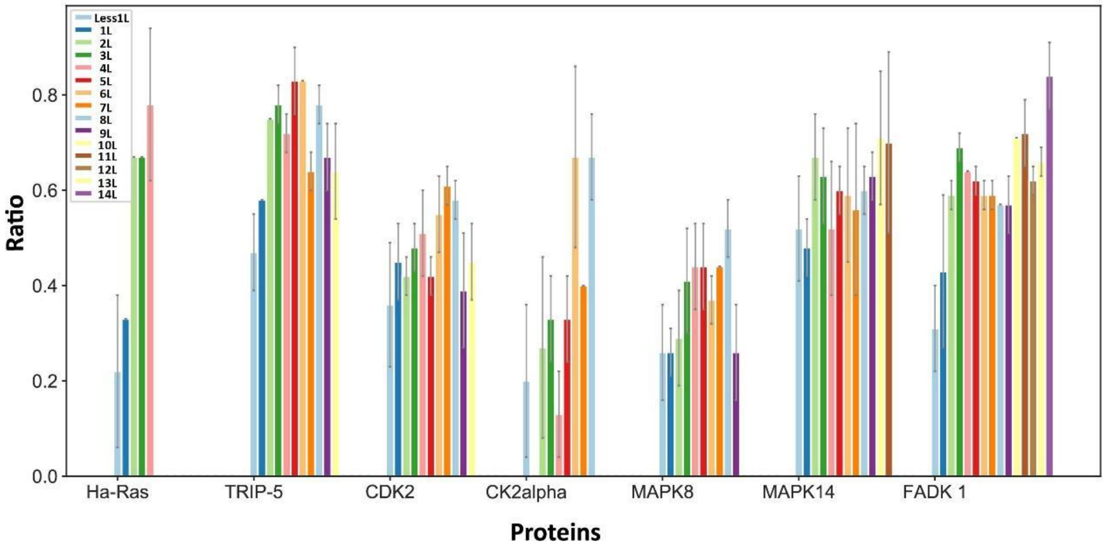


**Figure supplement 3**. Random sampling of homologous sequences. For each protein, we randomly sampled different numbers of homologous sequences such as 1L (L: length of protein), 2L, 3L and so on. The ratio refers to the proportion of identical key allo-residues identified by different numbers of homologous sequences and all homologous sequences.
